# Targeted control of gene expression using CRISPR-associated endoribonucleases

**DOI:** 10.1101/2025.03.17.643761

**Authors:** Sagar J. Parikh, Heather M. Terron, Luke A. Burgard, Derek S. Maranan, Dylan D. Butler, Abigail Wiseman, Frank M. LaFerla, Shelley Lane, Malcolm A. Leissring

## Abstract

CRISPR-associated endoribonucleases (Cas RNases) cleave single-stranded RNA in a highly sequence-specific manner, by recognizing and binding to short RNA sequences known as direct repeats (DRs). Here we investigate the potential of exploiting Cas RNases for the regulation of target genes with one or more DRs introduced into the 3’ untranslated region, an approach we refer to as DREDGE (<u>d</u>irect <u>r</u>epeat-<u>e</u>nabled <u>d</u>own-regulation of <u>g</u>ene <u>e</u>xpression). The DNase-dead version of Cas12a (dCas12a) was identified as the most efficient among 5 different Cas RNases tested and was subsequently evaluated in doxycycline-regulatable systems targeting either stably expressed fluorescent proteins or an endogenous gene. DREDGE performed superbly in stable cell lines, resulting in up to 90% downregulation with rapid onset, notably, in a fully reversible manner. Successful control of an endogenous gene with DREDGE was demonstrated in two formats, including one wherein both the DR and the transgene driving expression of dCas12a were introduced in one step by CRISPR-Cas. Our results establish DREDGE as an effective method for regulating gene expression in a targeted, highly selective, and fully reversible manner, with several advantages over existing technologies.

## 1. Introduction

CRISPR (clustered regularly interspaced short palindromic repeats) refers to a hallmark DNA sequence that plays a key role in antiviral defense in prokaryotes [1, 2]. Repeated elements, known as direct repeats (DRs), flank interspaced sequences, known as spacers, that are perfectly complementary to the sequences of viral genomes [3]. In response to viral infection, the CRISPR region is transcribed to generate a pre-crRNA, which is subsequently processed by CRISPR-associated (Cas) single-stranded endoribonucleases (RNases) to excise the spacer RNAs [2]. Cas RNases carry out pre-crRNA processing by recognizing and binding to cognate DRs in a highly sequence-specific manner, then cleaving the RNA within or adjacent to the DR sequence [2]. The excised spacer RNAs, also known as guide RNAs (gRNAs), in turn, associate with Cas double-stranded deoxyribonucleases (DNases) to target and cleave specific DNA sequences within the invading viral genome [1].

Relative to Cas DNases, which have been widely exploited for the development of sophisticated methods for modifying genomic DNA and other applications [4], Cas RNases have received considerably less attention, with only a small number of studies investigating their potential for gene regulation [5-10], and even then often as an adjunct to conventional CRISPR-Cas technology [11]. Our group has been developing and evaluating methods for achieving drug-inducible downregulation of genes, ultimately for in vivo applications that, in the ideal case, are completely reversible and perfectly selective. We previously developed a method predicated on doxycycline (Dox)-dependent histone methylation of the promoter region of a target gene, which successfully downregulates a target gene at very low concentrations of Dox and was found to be fully reversible in cell culture after one week of Dox exposure [12]. However, accruing evidence indicates that the Krüppel-associate box repressor (KRAB) domain used in this system can disrupt gene expression irreversibly in a subset of cells after prolonged exposure to Dox [13, 14, 15, 33], a possibility we explicitly aim to avoid, particularly in vivo.

We hypothesized that DRs and their cognate Cas RNases could be utilized as a powerful means for downregulating target genes selectively and fully reversibly, an approach we refer to as DREDGE (<u>d</u>irect repeat-<u>e</u>nabled <u>d</u>ownregulation of <u>g</u>ene <u>e</u>xpression). Specifically, by introducing one or more DRs into the transcribed portion of a target gene, a cognate Cas RNase can be directed to cleave the modified mRNA specifically and exclusively, leading to its subsequent destruction by any of several mechanisms. While multiple instantiations are possible, in this study we evaluate the approach of introducing one or more DRs into the 3’ untranslated region (UTR) of a target gene (Fig. 1A). In this approach, dubbed *3’ DREDGE*, a Cas RNase disrupts expression of the target gene by catalyzing the removal of the poly(A) tail from the target gene mRNA, which triggers its rapid destruction via deadenylation-dependent mRNA decay [16] (Fig. 1A). Relative to other approaches, there is a singular practical advantage to targeting the 3’ UTR: it allows for the possibility of introducing both the DR(s) (typically placed immediately downstream of the stop codon) <u>and</u> a transgene (TG) expressing the cognate RNase (inserted downstream of the 3’ end of the 3’ UTR) in one step via CRISPR-Cas—a significant time- and cost-saving measure, particularly for in vivo applications—which we implement herein.

**Figure 1.**
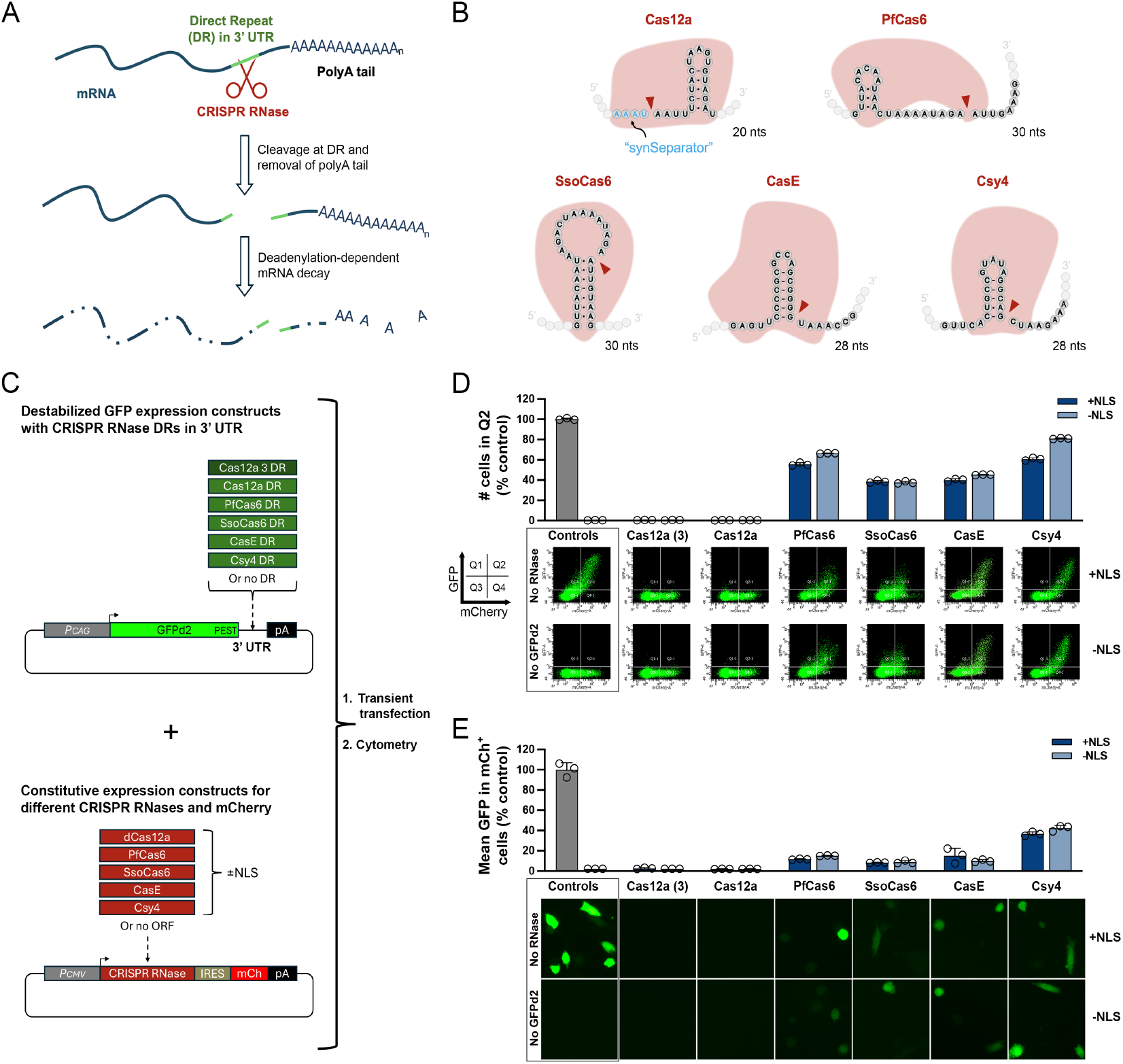
Screening of candidate Cas RNases for 3’ DREDGE. **A**, Mechanism of 3’ DREDGE. Cleavage of the DR(s) in the 3’ UTR of mRNA (green) by a Cas RNase (red) removes the poly(A) tail, triggering rapid degradation. **B**, DRs for the five Cas RNases investigated in this study, with cleavage sites indicated (red arrows). Note the “synSeparator” adjacent to the 5’ end of the Cas12a DR. **C**, Designs of constructs expressing GFPd2 with different DRs (or no DR) in the 3’ UTR (green) and constructs expressing individual Cas RNases (or no RNase) together with mCherry (red). **D**, Cell cytometry data for MEFs expressing different Cas RNases and GFPd2 constructs with their cognate DRs. Graph of the percentage of cells in Q2 relative to controls (top) within log-log plots of GFP vs. mCherry RFU values (n=3 replicates), with representative plots shown (bottom). **E**, Graph of GFP intensity in mCherry+ cells derived from RFU plots shown in **D** and normalized to controls, along with representative images of GFP fluorescence in cells prior to cytometry (bottom).

The present study evaluates for the first time, to our knowledge, the feasibility of exploiting the RNase activity of Cas12a (a.k.a., Cpf1)—specifically the DNase-dead version (dCas12a)—to downregulate the expression of target genes. This choice was inspired by a study showing that Cas12a completely eliminated the expression of a control protein whose transcript contained Cas12a DRs in its 3’ UTR [17]. A major goal of the latter study was to overcome this effect, so that Cas12a and its crRNA could be expressed on a single-transcript to, in one application, drive histone methylation via CRISPR interference (CRISPRi) using a single expression construct [17]. Ironically, the magnitude of the downregulation achieved with the latter approach in no case matched the essentially complete downregulation observed with the control vector featuring Cas12a DRs in its 3’ UTR [17], hence inspiring us to conceive of 3’ DREDGE.

We report here multiple successful implementations of 3’ DREDGE, including the fully reversible downregulation of the endogenous gene, *CTSD*, encoding cathepsin D (CatD), a lysosomal protease that our group has shown plays a critical role in the etiology of Alzheimer’s disease [18-20]. In a transient transfection paradigm, dCas12a and four different Cas RNases each markedly downregulated the expression of destabilized green fluorescent protein (GFPd2), with dCas12a markedly outperforming the other Cas RNases, achieving essentially 100% downregulation. In stable cell lines expressing GFPd2 constitutively and dCas12a in a Dox-dependent manner, DREDGE achieved >90% down-regulation with one or three Cas12 DRs and, moreover, exhibited full reversibility with a half-life (t_1/2_) of less than one day. We also utilized DREDGE to take control of the endogenous gene, *CTSD*, using CRISPR-Cas to introduce either three DRs alone or one DR together with a TG expressing dCas12a in a Dox-dependent manner. Both approaches resulted in efficient downregulation of *CTSD*—crucially—in a manner that was fully reversible after >2 months’ continuous exposure to Dox and also after repeated cycles of addition and withdrawal of Dox. Our results validate DREDGE as an effective means for downregulating target genes with full reversibility and—importantly—complete specificity, circumventing off-target effects that commonly plague other approaches such as RNA interference (RNAi) and CRISPRi [21, 22]. By virtue of these characteristics, DREDGE is expected to have broad utility for assessing biological responses to myriad transient phenomena ranging from injury, to toxin exposure, to drug treatments.

## 2. Materials and Methods

### 2.1. DNA constructs

All constructs were assembled using the HiFi DNA Assembly Kit according to manufacturer’s recommendations (New England Biolabs (NEB), Beverly, MA, USA) from DNA fragments generated either by restriction digestion (NEB, Beverly, MA, USA), by PCR with Q5^®^ DNA Polymerase (NEB, Beverly, MA, USA), or by de novo DNA synthesis (Integrated DNA Technologies (IDT), San Diego, CA, USA). All constructs were verified by next-generation DNA sequencing (Azenta Life Sciences, Waltham, MA, USA).

#### 2.1.1 Vectors for transient transfection experiments

The parent vector expressing destabilized GFP (GFPd2) under the control of the CAG promoter, pCAG-GFPd2 (Addgene plasmid #14760 [23]) was modified by inserting an open-reading frame (ORF) encoding a puromycin resistance cassette (Puro^r^)-T2A-TagBFP fusion protein (amplified from Addgene Plasmid #155307 [24]) into an AvrII restriction site between the SV40 promoter and SV40 poly(A) signal within the pCAG-GFPd2 vector. After sequence verification, de novo synthesized DNA sequences encoding one or more DRs for 5 different Cas RNases were inserted into a NotI site within the 3’ UTR. The ORFs for dCas12a without or with nuclear localization signals (NLSs) were generated by PCR from pSLQ10844 (Addgene Plasmid #183956 [25]). ORFs for the remaining Cas RNases (PfCas6, SsoCas6, CasE and Csy4) containing an N-terminal FLAG sequence (both without or with an adjacent NLS) were synthesized de novo, as mammalian codon-optimized gBlock DNA sequences based on amino acid sequences derived from Campa et al. [17]. The latter ORFs were inserted into the pICherryNeo vector (gift of Dario Vignali; Addgene Plasmid #52119) digested with XbaI.

#### 2.1.2 Vectors for inducible expression of Cas RNases (and mCherry and Neo^r^)

Vectors for inducible expression of Cas RNases (or no RNase) along with mCherry (mCh) with 3 C-terminal NLSs and neomycin/G418 resistance (Neo^r^) were generated as follows. A de novo synthesized DNA sequence encoding (1) appropriate restriction sites for in-frame insertion of ORFs for Cas RNases, (2) a P2A sequence, (3) mCh, (4) a T2A sequence, (5) Neo^r^, and (6) overlapping sequences was cloned into the EcoRI and NotI sites within pTet-One [26] (Takara Bio USA, Inc., San Jose, CA, USA) to create pMT_mCh_NeoR_pT1. The ORF for dCas12a (the codon-optimized version) containing 2 NLSs was amplified from pSLQ10875 (Addgene Plasmid #183962 [25]) and the ORFs for PfCas6, SsoCas6, CasE and Csy4 with NLSs were amplified from the pICherryNeo vectors described above and cloned into NcoI and SacI sites within pMT_mCh_NeoR_pT1.

#### 2.1.3. Targeting construct for inserting 3 Cas12a DRs into the 3’ UTR of the murine CTSD

HiFi DNA Assembly was used to assemble the following in pBlueScript KS II(+) (Agilent Technologies, Santa Clara, CA, USA) digested with NotI and KpnI: (1) a ∼750-bp 5’ homology arm containing overlap with the vector on the 5’ end, a LoxP site on the 3’ end, and modifications to the sequences within the *CTSD* ORF recognized by gRNAs used for insertion (see Supp. Fig. S2), PCR amplified from C57Bl/6J mouse tail DNA; (2) a PCR amplicon of the IRES from pICherryNeo with appropriate 5’ and 3’ overhangs; (3) the ORF for Puro^r^_P2A_GFP and a second LoxP site, amplified from Addgene Plasmid #111596 [27]; (4) a PCR amplicon of the 3 Cas12a DRs, amplified from that cloned into in pCAG-GFPd2; and (5) a ∼750-bp 3’ homology arm, with overlap for the pBlueScript vector, amplified from C57Bl/6J mouse tail DNA.

#### 2.1.4. Targeting construct for the DR+TG insert

The region encoding the artificial intron, the ORF for dCas12a, the IRES and the mCherry ORF was PCR amplified from the previously generated vector encoding dCas12a in pICherryNeo and cloned into pTet-One digested with EcoRI and NotI. The resulting vector, pdCas12a_pICNT, was digested with BamHI and NotI, and a PEST sequence, amplified from pCAG_GFPd2, was inserted in-frame into the C-terminus of dCas12a, to generate pdCas12a_PEST_pICNT. The MultiSite Gateway Pro kit was used to assemble the final construct, from 3 pieces combined into a fourth pDEST vector, according to manufacturer’s recommendations (Thermo Fisher Scientific, Waltham, MA, USA). The first piece was initially constructed in pBlueScript KS II(+) cut with SacI and XhoI, into which was inserted two de novo synthesized gBlocks encoding: (1) a synthetic gRNA (syn_gRNA) [28] for ultimate excision of the construct for CRISPR; (2) a ∼600 bp homology arm, with substitutions to remove the gRNA sequence used for modifying the *CTSD* allele; (3) a Cas12a DR flanked on the 5’ end by the synSeparator sequence (AAAT; [29]), positioned immediately downstream of the stop codon (see Supp. Fig. S3); (4) the endogenous *CTSD* 3’ UTR; and (5) an FRT sequence. Once assembled, the latter elements were amplified using primers containing appropriate attB sequences, then introduced into pDONR221 P1-P4 using BP Clonase II (Thermo Fisher Scientific, Waltham, MA, USA). The second piece was constructed by PCR amplifying a region encoding (1) the tetracycline response element (TRE); (2) the artificial intron; (3) the dCas12a with C-terminal PEST sequence; (4) the IRES; (5) the mCherry ORF; and (6) the poly(A) sequence, from pdCas12a_PEST_pICNT, using primers containing appropriate attB sequences, which was then introduced (in reverse orientation) into pDONR221 P4r-P3r. The third piece was generated by PCR amplifying, with primers containing appropriate attB sequences, a de novo synthesized gBlock encoding a ∼600-bp 3’ homology region downstream of the *CTSD* 3’ UTR followed by a second Syn_gRNA sequence. This sequence was introduced into pDONR221 P3-P2 using BP Clonase, then the CAG promoter, obtained from pCAG-GFPd2 as a SalI-SalI fragment was subsequently introduced (also in reverse orientation) into a SalI site within the latter gBlock. After sequence verification, the latter three vectors were assembled into pcDNA6.2™/V5-pL-DEST using LR Clonase II (Thermo Fisher Scientific, Waltham, MA, USA), generating a construct called pCTSD_DR+TG.

### 2.2. Cell culture

All experiments employed wild-type, SV40-immortalized mouse embryonic fibroblasts (MEFs; Cat #CRL-2907; ATCC, Manassas, VA, USA). Cells were maintained at 37 °C in a humidified incubator supplemented with 5% CO_2_ in DMEM containing Gluta-MAX® supplemented with 10% Tet System Approved Fetal Bovine Serum (Takara Bio USA, San Jose, CA, USA), 100U/mL penicillin, and 100 *μ*g/mL streptomycin.

### 2.3. Flow cytometry and FACS

For the screening of Cas RNases, MEFs were transfected with expression vectors using the Amaxa Nucleofector II electroporation system according to manufacturer’s recommendation (Lonza Bioscience, Bend, OR, USA) and cultured overnight. Cells were then imaged on a Zeiss Axiovert 200m fluorescent microscope (Zeiss, Dublin, CA, USA), harvested by trypsinization, and collected in tubes via a cell strainer. Flow cytometry on these cells and subsequently generated stable cell lines was performed on a BD LSRFortessa™ X-20 Cell Analyzer using lasers and filter sets optimized for the detection of GFP and mCherry fluorescence according to manufacturer’s recommendations (BD BioSciences, Franklin Lakes, NJ, USA). For the generation of stable cell lines, individual cells were plated into 96-well plates by fluorescence-activated cell sorting (FACS) using a BD FACSAria™ II Cell Sorter (BD BioSciences, Franklin Lakes, NJ, USA).

### 2.4. Targeted modification of the CTSD locus

Three Cas12a DRs were introduced into the endogenous *CTSD* locus of MEFs by transfecting cells as above with a mixture comprising (1) recombinant Alt-R A.s. Cas12a (Cpf1) *Ultra* Nuclease; (2) two synthetic gRNAs (gRNA_2 and gRNA_3; see Supp. Fig. S2B); and (3) 0.5 *μ*g of a PCR product of the targeted insert generated using phosphorothioate bond-containing oligonucleotides (IDT, San Diego, CA, USA). The DR+TG modification was done similarly, using Cpf1 Nuclease complexed to three synthetic gRNAs— gRNA_3, gRNA_9 (see Supp. Fig. S3B), and Syn_gRNA (GCTGTCCCCAGTGCATATTC) [28]—in this case using 2 *μ*g of pCTSD_DR+TG.

### 2.5. CatD Activity Assays

CatD activity assays were performed essentially as described [12, 18] using the internally quenched, fluorogenic peptide substrate (Mca-GKPILFFRLK(Dnp)-R-NH_2_) (InnoPep, Inc. San Diego, CA, USA). In a typical reaction, near-confluent monolayers of MEFs were detached by brief digestion with trypsin-EDTA. After 2 washes in PBS, pelleted cells were lysed in 200 *μ*L Lysis Buffer (40 mM NaOAc, 0.1% CHAPS, pH 3.5), incubated on ice for 10 min, then centrifuged at 21,000× g for 1 min. The concentration of the supernatant was estimated by A280 quantification using a NanoDrop® ND-1000 UV-Vis Spectrophotometer (Thermo Fisher Scientific, Waltham, MA, USA). Samples were loaded into black 384-well plates and reactions initiated by addition of an equal volume of Reaction Buffer (100 mM NaOAc, 0.2 M NaCl, pH 3.5) containing fluorogenic substrate (2 *μ*M). Plates were immediately loaded into a microplate reader (Gemini EM, Molecular Devices, LLC, San Jose, CA, USA), and fluorescence (λ_ex_ = 328 nm, λ_em_ = 393 nm) was read continuously every 15 s for ≥10 min. Proteolytic activity was determined from initial slopes of progress curves from each well, obtained using SoftMax Pro (v. 5.0; Molecular Devices, LLC, San Jose, CA, USA), normalized to individual A280 readings and appropriate controls.

## 3. Results

### 3.1. Selection and screening of candidate Cas RNases

To assess the feasibility of downregulating target genes using the 3’ DREDGE approach (Fig. 1A), we selected five different Cas RNases for investigation based on different criteria (Fig. 1B). Cas12a (a.k.a., Cpf1; Fig. 1B), our initial and primary candidate for reasons outlined above, is unusual in possessing both DNase and RNase activity; consequently, we utilized a DNase-dead version of the protein, specifically the engineered “hyperdCas12a” DNase-dead version from *Lachnospiraceae bacterium* developed by Guo and colleagues [25]. Further bolstering our assessment that Cas12a in particular might constitute an especially effective Cas RNase, Magnusson and colleagues designed a short A/U-rich “synthetic separator” (synSeparator; Fig. 1B)—AAAU—that enhances the excision of spacer sequences when positioned adjacent to the 5’ end of the Cas12a DR [29], which we incorporated into all Cas12a DR constructs (Fig. 1B). Two other Cas RNases, PfCas6 and SsoCas6 (Fig. 1B)—from *Pyrococcus furiosus* and *Sulfolobus solfataricus*, respectively—were selected based on reports that they are “multiple-turnover enzymes,” as distinct from single-turnover Cas RNases, which remain tightly bound to the cognate DR after cleavage [2]. Finally, CasE (a.k.a., EcoCas6e) and Csy4 (a.k.a., Cas6f)—from *Escherichia coli* and *Pseudomonas aeruginosa*, respectively—were selected based on a study showing these to be the best-performing Cas RNases among 9 tested when introduced into mRNAs [17].

As illustrated in Figure 1B, the cognate DRs of the five Cas RNases contain certain commonalities and some key differences [11, 30-32]. The DRs are all quite short, comprising ≤30 nucleotides, but highly varied in primary nucleotide sequence (Fig. 1B). The DR for Cas12a is the shortest, comprising just 20 nucleotides (neglecting the four-nucleotide synSeparator included in all constructs) (Fig. 1B). A second distinguishing feature is the placement of the cleavage sites. All the tested Cas RNases except Cas12a cleave within the DR, at a position seven or eight nucleotides from the 3’ end; Cas12a, by contrast, cleaves outside the DR, at the 5’ end (Fig. 1B). Finally, the five DRs differ in the extent to which they form hairpins, and the degree of hydrogen bonding within each hairpin, with CasE featuring a hairpin comprised of six G/C base pairs [11], and PfCas6 featuring a hairpin comprised of just three A/U base pairs [30].

To compare the relative efficacy of the five selected Cas RNases for downregulating a target gene, we generated a two-part model system that uses GFP fluorescence as a convenient marker of target gene expression, and mCherry as a marker of Cas RNase expression (Fig. 1C). The first part of this system consisted of vectors expressing a destabilized form of GFP (GFPd2), featuring a very short, two-hour half-life (t_1/2_) [23], driven by the strong CAG promoter. We cloned one DR for each of the five Cas RNases into the 3’ UTR of this construct, or no DR as a control (Fig. 1C, top). For dCas12a, we also tested a three-DR version, wherein the DRs flank two “dummy” spacers with no complementarity to mouse genomic DNA [29] (Fig. 1C, top; see Fig. 4B). For the second part of this system, each of the five Cas RNases was cloned into a vector co-expressing mCherry (Fig. 1C, bottom). For each Cas RNase, we generated versions either lacking or containing a nuclear localization signal (NLS) (two in the case of Cas12a), for a total of 10 Cas RNase expression constructs plus an empty-vector, no-RNase control (Fig. 1C).

Each Cas RNase/mCherry expression construct was cotransfected together with the GFPd2 construct bearing the cognate DR(s) in its 3’ UTR into mouse embryonic fibroblasts (MEFs). Controls consisted of: (1) MEFs transfected with empty (No-RNase) mCherry vector plus GFPd2 lacking a DR (No-DR), representing the maximum possible GFP fluorescence; and (2) cells transfected with empty mCherry vector alone (No-GFPd2), representing the minimum possible GFP fluorescence (Fig. 1D). One day after transfection, cells were harvested and analyzed by FACS. Log-log plots of the fluorescence in each cell were generated, depicting GFP and mCherry on the Y- and X-axes, respectively, which were further divided into 4 quadrants, by using untransfected MEFs as a non-fluorescent control to establish strict boundaries for GFP and mCherry fluorescence (Fig. 1D). In control cells cotransfected with No-DR GFPd2 and No-RNase mCherry, abundant GFP and mCherry fluorescence were present, resulting in large numbers of cells appearing in the upper-right quadrant (Q2) of RFU plots; conversely, for cells expressing mCherry alone, no cells were present in Q2, as expected (Fig. 1D). In marked contrast, cells expressing each of the five Cas RNases and their cognate DRs all showed decreases in the percentage of cells in Q2 to varying extents relative to No RNase controls (Fig. 1D), together with substantial reductions in the mean level of GFP fluorescence in mCherry-positive cells (e.g., cells in Q2 and Q4; Fig. 1E). Significantly, dCas12a performed superiorly relative to all other Cas RNases tested by both metrics, with almost no cells appearing in Q2 (Fig. 1D) and GFP fluorescence levels being indistinguishable from No-GFPd2 controls (Fig. 1E). There were no detectable differences between GFPd2 constructs expressing one or three Cas12a DRs and dCas12a in this experimental paradigm (Fig. 1D,E).

The absence or presence of an NLS on the Cas RNases only modestly impacted their performance (Fig. 1D, E). Marginal decreases in efficacy were observed for a few Cas RNases lacking an NLS, although for others the NLS had either no effect or the opposite effect (Fig. 1D, E). Nevertheless, this parameter was sufficiently important to us to warrant additional testing. Consequently, we generated an additional dCas12a/mCherry expression construct containing a nuclear exclusion sequence (NES), which was compared in parallel with the dCas12a constructs lacking or possessing two NLS sequences in cells coexpressing GFPd2 with one Cas12a DR (or no DR) in its 3’ UTR. Consistent with the previous results, no significant differences were observed between the constructs containing an NES, an NLS or neither localization sequence (Supp. Fig. S1). Nevertheless, in light of the modest improvements in performance observed for some RNases possessing NLSs, we elected to proceed with dCas12a constructs containing two NLS sequences.

### 3.2. Dox-regulatable gene expression by 3’ DREDGE using dCas12a

We next sought to characterize the performance of cells expressing dCas12a in a doxycycline (Dox)-dependent manner, using the Tet-One™ system [26] (Takara Bio USA, Inc., San Jose, CA, USA). To that end, we first created cell lines stably expressing GFPd2 with zero, one or three Cas12a DRs (Fig. 2A). These stable cell lines were subsequently used to generate double-stable lines also expressing dCas12a (with two NLSs) together with mCherry (with three NLSs) and neomycin resistance, all under the control of the TRE promoter to permit Dox-regulatable expression. The latter construct lacking dCas12a served as a No-RNase control. As illustrated in Figure 2C and D, all control lines performed as anticipated. For the No-DR GFPd2 cells lines, essentially 0% and 100% of cells appeared in Q2 in the absence or presence of Dox, respectively, whether expressing dCas12a or no RNase (Fig. 2C, left), and mean GFP levels were essentially unchanged, irrespective of RNase expression or Dox administration (Fig. 1D, left). Cells stably expressing GFPd2 with either one or three Cas12a DRs in the 3’ UTR behaved similarly when co-expressing the No-RNase Dox-inducible control vector (Fig. 1C,D, middle and right). In striking contrast, expression of dCas12a in GFPd2 lines with one or three DRs elicited marked reductions in the number of cells in Q2 (89.4% and 84.8%, respectively) relative to the same parental lines expressing No-RNase mCherry controls (Fig. 2C, middle and right). Similarly, relative to cells without Dox treatment, cells with Dox-induced dCas12a expression exhibited 93.2% and 93.7% reductions in mean GFP fluorescence in lines with one and three Cas12a DRs, respectively (Fig. 2D, middle and right).

**Figure 2.**
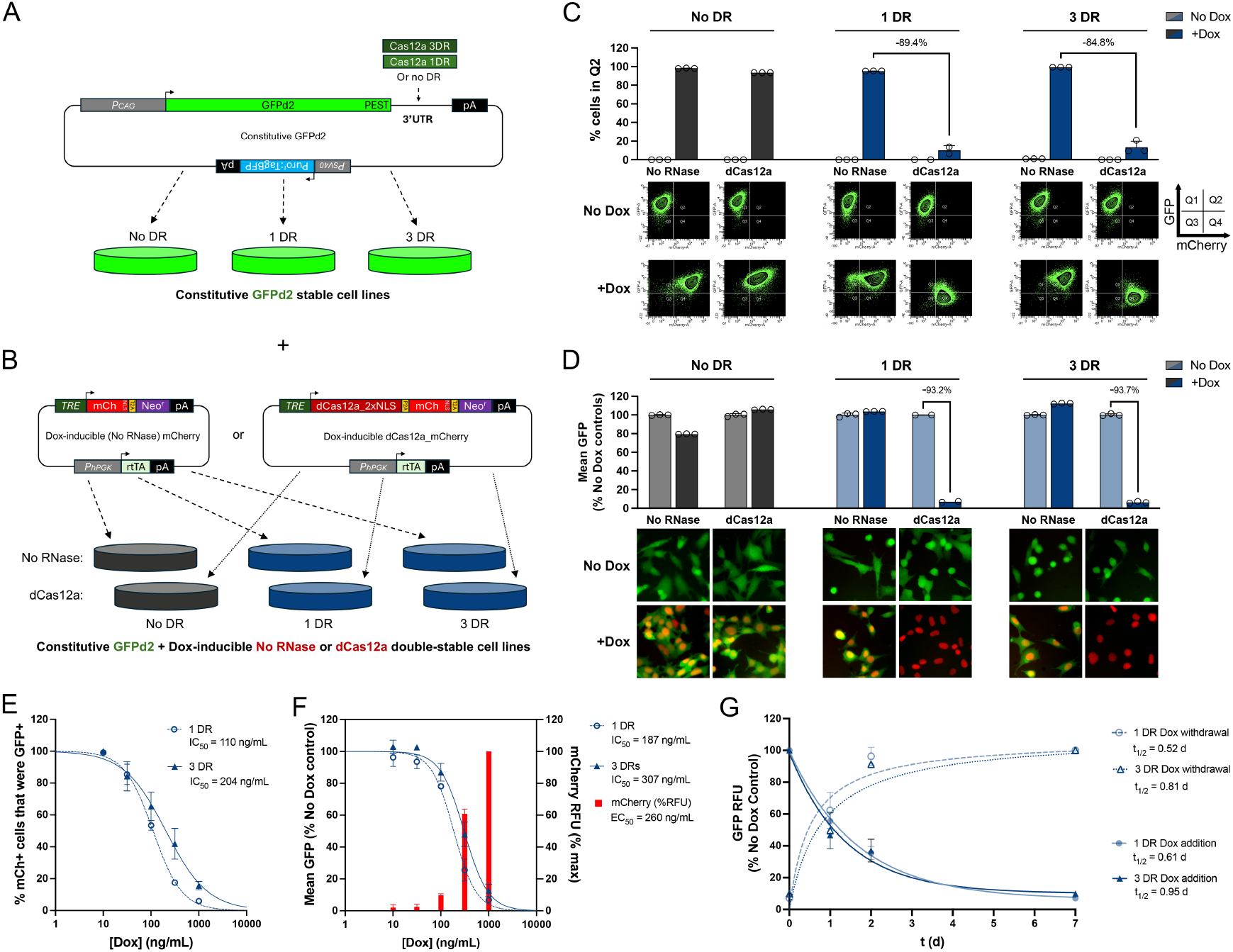
Dox-regulatable control of gene expression by 3’ DREDGE. **A**, Design of constructs constitutively expressing GFPd2 with zero, one, or three Cas12a DRs in the 3’ UTR used to create three different stable cell lines. **B**, Design of constructs with Dox-regulatable co-expression of dCas12a (or No RNase) and mCherry used to create double-stable cell lines from the lines in **A. C**, Percentage of cells in Q2 for double-stable cell lines expressing GFPd2 with zero, one, or three DRs and also conditionally expressing either Cas12a or no RNase, tested in the absence or presence of Dox (top) derived from log-log plots of GFP vs. mCherry RFU (bottom). **D**, Mean GFP RFU in the cell lines in **C** in the absence or presence of Dox derived from cell cytometry (top) with representative images of cells in the different conditions (bottom). Data in **C** and **D** are normalized to Dox-treated No-DR and No-RNase controls; n=2-3 per condition. **E**,**F**, Dose-response curves showing (**E**) the percent of mCherry+ cells (i.e., Q2+Q4) that were also GFP+ (i.e., in Q2) and (**F**) mean GFP RFU as a function of Dox dose in stable cell lines with one or three DRs conditionally expressing cDas12a. Red columns in **F** show the mean mCherry RFU as a function of Dox dose for both cell lines, normalized to the maximum at 1000 ng/mL. Mean IC50 values for all dose-responses are indicated. Data are mean ± SEM, normalized to values in the absence of Dox for each line; n=2-3 per condition. **G**, Time courses of GFP RFU in response to addition (solid lines) or withdrawal (dashed lines) of Dox in the cell lines in **E** and **F** normalized to No-Dox controls. Mean half-life values (t1/2) are shown. Data are mean ± SEM for 2-3 independent experiments.

To more completely characterize the performance of 3’ DREDGE we used these stable lines to carry out Dox dose-response experiments (Fig. 2E,F) as well as time courses of the responsiveness of gene expression after addition or removal of Dox (Fig. 2G). Dose-response experiments conducted in the cell lines with one and three DRs revealed IC_50_s of 110 and 204 ng/mL Dox, respectively, using the percent of mCherry-positive cells (Q2+Q4) present in Q2 as a metric (Fig. 2E). Similar results were obtained using mean GFP fluorescence in all cells, yielding IC_50_s of 187 and 307 ng/mL Dox, respectively (Fig. 2F). Notably, the mean mCherry fluorescence (representing the average in all runs in both cell lines) exhibited a similar EC_50_ of 260 ng/mL Dox (Fig. 2F), suggesting that the percent downregulation of GFPd2 was essentially a direct reflection of dCas12a expression. Significantly, time courses revealed that the DREDGE approach exhibits remarkably fast kinetics, with t_1/2_s for induction of downregulation of 0.52 and 0.81 d for one and three DRs, respectively (Fig. 2G). The t_1/2_s for restoration of activity were similar: 0.61 and 0.95 d, respectively. Overall, both the one-DR and the three-DR systems performed comparably; however, it is noteworthy that the one-DR system consistently performed marginally better on all the foregoing measures (see *Discussion*).

### 3.3. Dox-regulatable expression of the endogenous gene, CTSD, by 3’ DREDGE

We next sought to assess the feasibility of downregulating an endogenous gene with the 3’ DREDGE approach. Having previously developed an alternative method for down-regulating *CTSD* [12], we selected this gene as our target. We tested two basic configurations (Figs. 3 and 4). In the first, we used CRISPR-Cpf1 in MEFs to introduce three Cas12a DRs within the 3’ UTR of *CTSD* (Fig. 3A; Supp. Fig. S2). To this end, we used a gene-trap approach with a knockin construct featuring an internal ribosomal entry site (IRES) driving expression of a puromycin resistance gene (Puro^r^) fused to GFP, both flanked by LoxP sites, upstream of three Cas12a DRs, all flanked by ∼600-bp 5’ and 3’ homology arms (Fig. 3A; Supp. Fig. S2A). This construct lacked a promoter and also a poly(A) tail, so it could express Puro^r^ if and only if it integrated successfully into the 3’ UTR of *CTSD*. After puromycin selection, we identified individual positive clones by PCR, including one line featuring one copy of the integrated construct and one *CTSD* allele inactivated by NHEJ (Supp. Fig. S2D), which was used for all downstream analyses. After removal of the floxed IRES_ Puro^r^::GFP elements with Cre-recombinase, the modified CTSD allele contained only the three Cas12a DRs (plus an upstream LoxP site) (Fig. 3B; Supp. Fig. S2A).

**Figure 3.**
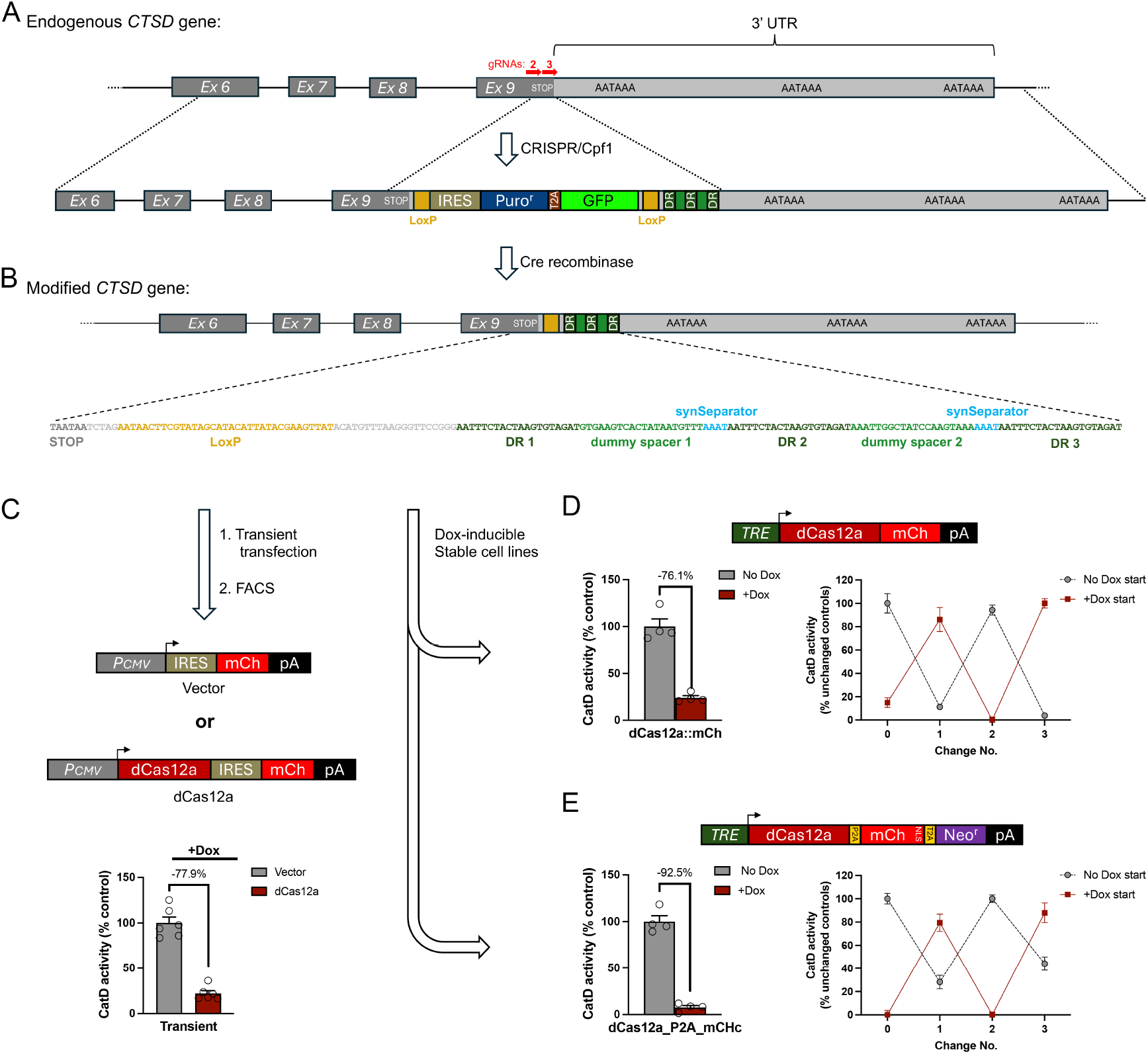
Dox-regulatable control of endogenous *CTSD* expression by 3’ DREDGE. **A**, Genomic structure of the 3’ end of murine *CTSD* (top) and design of the “gene-trap” targeting construct used to insert 3 Cas12a DRs into the 3’ UTR by CRIPSR/Cpf1 (bottom). Note the absence of a poly(A) signal within the targeting construct, and the presence of an upstream IRES and LoxP sites flanking all of the inserted region except the 3 Cas12a DRs. **B**, Design of the modified *CTSD* gene after targeted insertion of the gene-trap construct and removal of the IRES and Puro^r^-T2A-GFP ORF by Crerecombinase. The sequence of the inserted region, beginning at the stop codon in Exon 9 is shown. **C**, Overview of the design of DNA constructs used for transient transfection experiments (top) and the outcome of CatD activity assays performed on mCherry+ cells collected 24 h later by FACS (bottom). Data are mean ± SEM; n = 6. **D**,**E**, Stable cell lines created with a 3DR-containing parental cell line based on two Dox-regulatable constructs expressing a dCas12a::mCherry fusion (**D**, top) or the construct described in Fig. 2B, co-expressing dCas12a, mCherry and Neo^r^ from a single transcript via 2A peptides (**E**, top). Shown are Dox-dependent downregulation of CatD proteolytic activity achieved in the initial characterization both constructs (**D**,**E**, bottom left) and after repeated changes from +Dox to no Dox and vice versa spaced ∼1 week apart (**D**,**E**, bottom right). Data are mean ± SEM; n = 4-8 replicates per condition.

**Figure 4.**
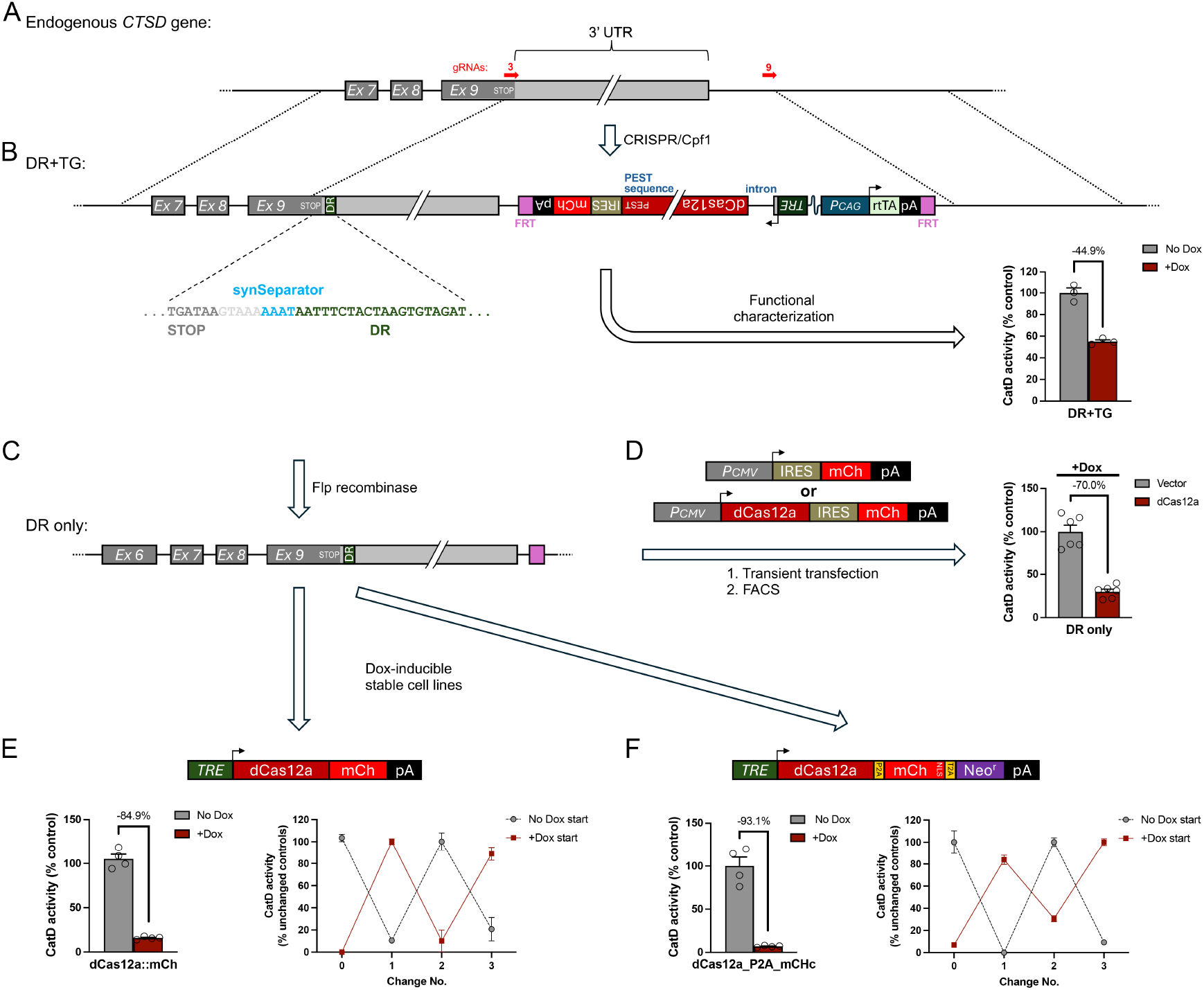
Dox-regulated control of endogenous *CTSD* expression using the “DR+TG” approach. **A**, Genomic structure of the 3’ end of murine *CTSD*, indicating the locations of gRNAs used for CRISPR/Cpf1. **B**, Design of the DR+TG insert used to introduce both a single dCas12a DR immediately downstream of the stop codon (sequence shown) and a complete transgene (TG) for Dox-regulatable expression of dCas12a flanked by FRT sites (purple). Note the inclusion of several design elements subsequently deemed to be suboptimal, including an artificial intron after the TRE and a C-terminal PEST sequence on dCas12a. Functional characterization of a line with one copy of the modified allele and one allele functionally knocked out by NHEJ (bottom right). Note that CatD activity was decreased by only ∼45%. Data are mean ± SEM; n = 3 replicates per condition. **C**, Design of the “DR-only” modified allele, after removal of the TG with Flp-recombinase. **D**, Design of DNA constructs used for transient transfection experiments (left) and the outcome of CatD activity assays performed on mCherry+ cells collected 24 h later by FACS (right). Data are mean ± SEM; n = 6. Note that larger reductions in CatD activity were achieved by this method. **E**,**F**, Design and performance of stable cell lines generated from the DR-only cell line based on Dox-regulatable constructs expressing a dCas12a::mCherry fusion (**E**, top) or co-expressing dCas12a, mCherry, and Neo^r^ (**F**, top). Shown are Dox-dependent downregulation of CatD activity achieved in the initial characterization both constructs (**E**,**F**, bottom left) and after repeated changes from +Dox to no Dox and vice versa spaced ∼1 week apart (**E**,**F**, bottom right). Data are mean ± SEM; n = 4-8 replicates per condition.

To assess the ability of this modified allele to effect the downregulation of CatD, we transiently transfected this cell line with either a constitutive dCas12a/mCherry expression vector or an empty (No-RNase) mCherry-only vector (Fig. 3C). One day later, we collected mCherry-positive cells from both conditions by FACS, and conducted CatD activity assays (see *Materials and Methods*), which revealed that CatD levels in dCas12a-expressing cells were reduced by 77.9% relative to cells transfected with the No RNase control (Fig. 3C).

To generate stable clones, we transfected the modified three-DR cell line with Dox-inducible vectors expressing dCas12a of two different designs, both co-expressing mCherry either as a fusion with dCas12a (Fig. 3D) or as a separate protein (via the incorporation of a P2A element; Fig. 3E). After addition of Dox to induce dCas12a expression, these stable cell lines exhibited marked reductions in CatD activity—of 76.1% and 92.5%, respectively—relative to the same cell lines maintained in the absence of Dox (Fig. 3D, E, left). Importantly, CatD activity within these lines in the absence of Dox was comparable to control lines expressing No-RNase control vectors, whether treated with Dox or not, even after extensive incubation with Dox (for ∼2 months) during the selection of these lines. Moreover, the two dCas12a-expressing lines both exhibited full reversibility even after being subjected to multiple successive alternating treatments without or with Dox (Fig. 3D, E, right).

In the second configuration, we aimed to introduce <u>both</u> a single Cas12a DR within the 3’ UTR of *CTSD* (immediately downstream of the stop codon) <u>and</u> a complete transgene (TG) for Dox-inducible dCas12a expression (downstream of the 3’ end of the *CTSD* 3’UTR), a configuration we refer to as “DR+TG” (Fig. 4A, B). Note that we included two FRT sites flanking the TG portion of the insert, so that it could be removed by Flp recombinase if desired (Fig. 4A). As before, CRISPR-Cpf1 was used introduce the DR+TG construct into the endogenous *CTSD* gene, in this case using two gRNAs, one near the stop codon of the *CTSD* ORF and the other downstream of the 3’ end of the *CTSD* 3’ UTR (Fig. 4A; Supp. Fig. S3). For this construct, we also incorporated a third “synthetic gRNA” (Syn-gRNA) sequence outside of each ∼600-bp homology arm, which enabled the DR+TG construct to be excised from the vector backbone simultaneously with CRISPR-Cas-mediated recombination, a technique shown to improve CRISPR efficiency [28]. Despite the presence of significant homology in the middle of our construct (∼700 bp)—in the form of the complete 3’ UTR of CTSD (plus one DR)—we encountered no difficulty identifying a clone with one copy of the DR+TG construct successfully integrated (and the other allele inactivated by NHEJ) (Supp. Fig. S3).

When CatD activity within this DR+TG cell line was quantified in the absence vs. the presence of Dox, CatD activity was significantly reduced by Dox, albeit only by 44.9% (Fig. 4B, right). There are several different plausible explanations for this less-than-ideal outcome, ranging from the fundamental to the technical, and it was important to distinguish among them. On the fundamental side, it could be that a single copy of the dCas12a TG is simply insufficient to effectively downregulate CatD using the DREDGE method. Alternatively, perhaps a single DR might be insufficient to effect complete downregulation irrespective of dCas12a levels. On the technical side, it could be that an active *CTSD* allele might be present that escaped our detection or was operative despite the NHEJ deletion detected by PCR. Another plausible technical explanation pertained to the design of the Dox-inducible dCas12a TG. The DR+TG construct was designed and created early on in this study, well before we had the benefit of experience with alternative dCas12a transgene designs, and it contained a few sub-optimal features, including (1) a PEST sequence at the C-terminus of dCas12a (which we had introduced to promote rapid recovery of CatD activity after Dox withdrawal) (Fig. 4B), (2) an intron after the TRE (which we had postulated would be beneficial for in vivo applications to promote mRNA processing) (Fig. 4B), and (3) an ORF for dCas12a that was not codon optimized to ensure no cryptic splice sites or polyadenylation signals were present.

To discriminate among these possibilities, we removed the dCas12a TG portion within the DR+TG line using Flp recombinase (Fig. 4C; Supp. Fig. S3). This “DR-only” line was then transiently transfected with a vector constitutively expressing dCas12a and mCherry (or mCherry-only vector) and, after collection of mCherry-positive cells from both conditions, CatD activity was assessed in the presence of Dox (Fig. 5D). CatD activity was decreased more completely in this paradigm (by 70.0%), suggesting that there was in fact a single functional *CTSD* allele in this line that could, in fact, be downregulated more completely (Fig. 5D). To confirm this, we generated stable cell lines expressing the two versions of Dox-inducible dCas12a expression vectors (or No-RNase controls) tested previously in the three-DR cell line (Fig. 4E, F). These cell lines behaved similarly to the three-DR stable lines, with dCas12a-expressing cells showing marked decreases in CatD activity of 84.9% and 93.1%, respectively, relative to the same lines grown in the absence of Dox. Moreover, as was the case for the three-DR cell lines, downregulation of CatD activity by both expression vectors in this single-DR line was fully reversible following multiple rounds of alternation between Dox addition and withdrawal (Fig. 4E, F, right panels).

## 4. Discussion

Our results establish DREDGE as an attractive general approach for regulating gene expression with several distinct advantages over widely used alternative methods. First, DREDGE offers a *high degree of selectivity* for the targeted gene, by virtue of the inclusion of one or more DRs as a *cis* element within the targeted gene. Both RNAi and CRISPRi, by contrast, are known to have significant off-target effects that are difficult to predict and control for, owing to their reliance on RNA::RNA and RNA::DNA complementarity, respectively, which tolerate mismatches to varying degrees [21, 22]. Second, as we show, DREDGE can be used to control gene expression in a *completely reversible* manner, a distinct advantage vis-à-vis CRISPRi and other approaches dependent upon histone methylation, which have been documented to result in irreversible methylation in several systems [13-15, 33]. Third, DREDGE controls gene expression with very *rapid kinetics*, both for downregulation, due to the high efficiency of deadenylation-dependent mRNA decay [16], and for recovery therefrom, as a consequence of targeting a constantly replenished pool of mRNA. These kinetic properties, in particular, should facilitate the study of the consequences of transient disruptions to gene expression, as can occur, for example, by exposure to discrete toxins [34]. Fourth, as implemented here in multiple configurations, DREDGE consistently resulted in a *higher degree of downregulation* (>80%) than is typical for RNAi and CRISPRi (<70%) [17, 25, 35], which can facilitate the study of phenotypes dependent upon relatively complete downregulation. Finally, DREDGE requires *minimal modifications to the target gene* (e.g., as small as 20 nucleotides in the case of the DR for Cas12a), which compares favorably to the size of other *cis* elements utilized for reversible gene regulation (e.g., >250 nucleotides in the case of the TRE [26]). The very small size of DRs minimizes the chances of perturbing expression of the targeted gene by affecting mRNA secondary structure or disrupting possible regulatory elements, such as microRNA binding sites [36].

The 3’ DREDGE approach specifically evaluated in this study has further unique advantages vis-à-vis alternative implementations. For instance, targeting the 3’ UTR makes it possible to regulate genes with multiple isoforms due to alternative transcription start sites, because such genes frequently share a common stop codon [37]. Moreover, as we demonstrated, targeting the 3’ UTR also makes it feasible to introduce both the DR and a transgene for a Cas RNase in one step using CRISPR-Cas. This feature of 3’ DREDGE makes it especially attractive for implementation in animal models, where single-locus modifications for gene regulation are strongly favored over the alternative of crossing lines with multiple modified genetic loci.

Our study also reveals potential limitations of 3’ DREDGE, at least for the specific implementation tested herein. In particular, continuous, very high levels of expression of Cas RNases appear to be required to efficiently downregulate target genes. In the example of our DR+TG approach, where just a single copy of the transgene was integrated, we obtained only partial (∼45%) downregulation. Although our results suggest that these less-than-ideal results may be attributable to suboptimal features of the transgene design, they establish the point that, as implemented here with dCas12a, the efficiency of DREDGE is directly determined by Cas RNase expression levels. Since the expression of endogenous genes can vary greatly, DREDGE may not be suitable for genes expressed at particularly high levels (unless further optimized; see below). Fortunately, there are resources that estimate the numbers of mRNA molecules in different cell-types and tissues [38], which may help assess the a priori suitability of DREDGE for specific genes.

To our knowledge, this is the first study to investigate the potential of Cas12a for regulating gene expression via its RNase activity. Recent studies offer insights into the RNase activity of Cas12a that are highly relevant to its performance in DREDGE. Detailed kinetic analyses reveal that Cas12a interacts with the pre-crRNA via exquisitely potent interactions, with equilibrium constants in the range of 56 pM to 3 nM for the RNA::protein complex, depending upon the species studied [39-41]. Given that RNA binding is the rate-limiting step in the processing of mRNAs by Cas12a [41], this is a favorable property. On the other hand, the half-life for dissociation of the Cas12a::pre-crRNA complex was recently reported to be 26 h [41], indicating that RNA binding is essentially *irreversible* and implying that only one mRNA molecule can be degraded by each Cas12a molecule. Fortunately, Cas12a cuts at the 5’ end of the DR (see Fig. 1B), so it remains associated with the 3’ end of the mRNA and thus does not interfere with deadenylation-dependent decay of the remainder of the mRNA. However, the essentially irreversible binding of Cas12a to each cognate DR suggests that Cas12a might be more effective if implemented with a single DR rather than multiple DRs, a prediction that is supported by the results we obtained here for systems with one vs. three DRs (see Fig. 2E,F).

Although Cas12a performed well in 3’ DREDGE, certain features of Cas12a could potentially be optimized to further improve its performance, though significant additional work would be required. For example, because it possesses both DNase and RNase activities, Cas12a is an especially large protein (∼140 kDa); expression might be made more efficient if truncated versions of the protein containing only the RNase activity could be developed. Also, it seems conceivable that the RNA::protein interaction could plausibly be engineered to render this interaction reversible and thereby convert Cas12a to multiple-turnover Cas RNase[2]; this might be achieved by varying the sequence of the DR and/or by manipulating resides within Cas12a involved in DR binding.

## 5. Conclusions

We conclude that DREDGE is an effective general method for regulating endogenous gene expression, particularly in circumstances where a high degree of selectivity and full reversibility are paramount, such as animal modeling studies. The 3’ DREDGE approach in particular also allows for a convenient method for rapid implementation of this system in one step via CRISPR-Cas, using the DR+TG approach; however, our results also illustrate that the effectiveness of Cas RNases expressed via a single copy of a transgene depends critically upon the relative expression level of the target gene and a suitably effective transgene design. Although our results establish dCas12a as an efficient mediator of 3’ DREDGE, the particular instantiation of this approach tested here is unlikely to be the most efficient of all possible configurations. The general approach might be improved by further engineering of dCas12a or, potentially, by discovery of alternative Cas RNases with higher RNA processing efficiency. We hope our findings will stimulate further improvements to this fundamental approach for targeted control of gene expression.

## Supplementary Materials

The following supporting information Is associated with this manuscript. Figure S1: Effects of nuclear localization and nuclear exclusion signals on the performance of 3’ DREDGE using dCas12a; Figure S2: Overview of genotyping results confirming the successful introduction of 3 Cas12a DRs into the 3’ UTR of murine *CTSD* via CRISPR-Cas; Figure S3: Overview of genotyping results confirming the successful introduction of the DR+TG insert into the 3’ region of murine *CTSD* via CRISPR-Cas.

## Author Contributions

Conceptualization, M.A.L.; methodology, M.A.L.; validation, S.J.P., H.M.T., L.A.B., D.S.M., D.D.B., A.W., S.L. and M.A.L.; data analysis, S.J.P., H.M.T., S.L. and M.A.L.; investigation, S.J.P., H.M.T., L.A.B., D.S.M., D.D.B., A.W., S.L. and M.A.L.; resources, F.M.L. and M.A.L.; writing—original draft preparation, M.A.L.; writing—review and editing, S.J.P. and S.L.; supervision, F.M.L., S.L. and M.A.L.; funding acquisition, F.M.L. and M.A.L. All authors have read and agreed to the published version of the manuscript.

## Institutional Review Board Statement

Not applicable.

## Informed Consent Statement

Not applicable.

## Data Availability Statement

Data are contained within the article and supplementary material. Original, raw fluorescence microscopy image files are deposited at the Open Science Framework repository, available at: www.doi.org/10.17605/OSF.IO/KUMJR.

## Acknowledgments

The authors thank the laboratory of Dr. Jorge Busciglio for generously providing access to their multiplate reader. We are grateful to Pauline Nguyen for expert assistance with cell cytometry and FACS analysis within the UCI Stem Cell Research Center. We thank Connie Cepko, Stanley L. Qi, Luke A. Gilbert, and Dario Vignali for providing cloning vectors via Addgene.

