## Supplemental Information for "Targeted control of gene expression using CRISPR-associated endoribonucleases"

Supplemental Materials
for

### Targeted control of gene expression using CRISPR-associated endoribonucleases

**Sagar J. Parikh[1], Heather M. Terron[1], Luke A. Burgard[1,2], Derek S. Maranan[1,2], Dylan D. Butler[1,2], Abigail Wiseman[1], Frank M. LaFerla[1,2], Shelley Lane[1] and Malcolm A. Leissring[1,*]**

[1] Institute for Memory Impairments and Neurological Disorders, University of California, Irvine, Irvine, CA 92697, USA
[2] Department of Neurobiology and Behavior, University of California, Irvine, Irvine, CA 92697, USA

#### Contents

| pp. | Fig./Table | Title |
|---|---|---|
| 2 | Fig. S1 | Effects of nuclear localization and nuclear exclusion signals on the performance of 3' DREDGE using dCas12a. |
| 3 | Fig. S2 | Overview of genotyping results confirming the successful introduction of 3 Cas12a DRs into the 3' UTR of murine *CTSD* via CRISPR-Cas. |
| 4 | Fig. S3 | Overview of genotyping results confirming the successful introduction of the DR+TG insert into the 3' region of murine *CTSD* via CRISPR-Cas. |

**Supplementary Figure S1.**

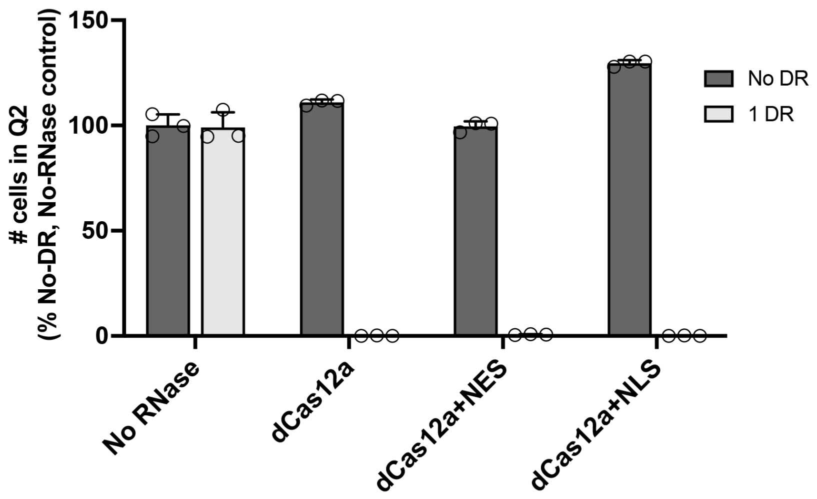


**Figure S1.** Effects of nuclear localization and nuclear exclusion signals on the performance of 3' DREDGE using dCas12a. Graph of the percentage of cells in Q2 (representing mCherry+ cells that are also GFP+) in MEFs transiently co-transfected with pCAG-GFPd2 expression vectors containing either 0 (No DR) or 1 DR, together with vectors expressing mCherry alone (No RNase) or together with dCas12a with no nuclear localization or exclusion signals (dCas12a), dCas12a with a nuclear exclusion sequence (dCas12a+NES), or dCas12a with 2 nuclear localization sequences (dCas12a+NLS). All data were normalized to cells transiently co-transfected with No-DR GFPd2 and No-RNAse mCherry vectors. Data are mean ± SEM for 3 replications per condition.

#### Supplementary Figure S2.

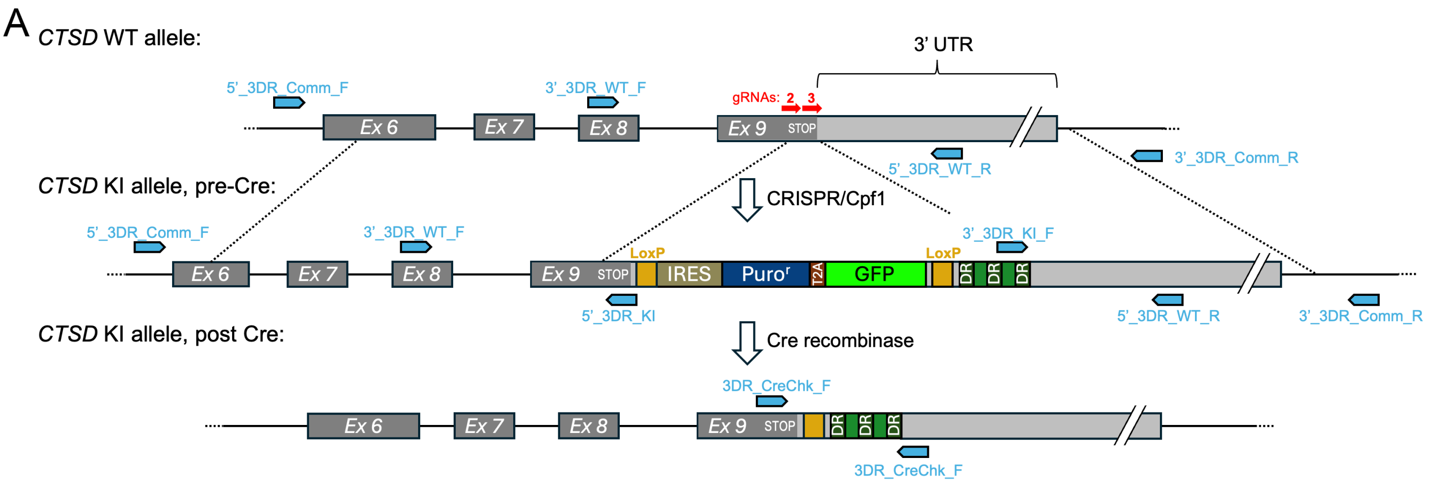


B gRNA design and alterations to knock-in insert:

| | gRNA_2 | gRNA_3 |
|---|---|---|
| gRNA sequence | ACAGAGACAACAATAGGGTCG | CCAATGCTGTCGTACTCTAAC |
| Endog. sequence | ...TTTGACAGAGACAACAATAGGGTCGGC... | ...TTTGCCAATGCTGTCGTACTCTAACTT... |
| A.A. sequence | PheAspArgAspAsnAsnArgValGly | PheAlaAsnAlaValValLeuSTP |
| KI insert sequence | ...TTCGATAGGGACAACAACAGAGTGGGC... | ...TTCGCGAACGCCGTAGTCCTGTGATAA... |

C Genotyping primers:

| Primer Name | Primer Sequence |
|---|---|
| 5'_3DR_Comm_F | CTGATCACTTACTTCCTCCTAGG |
| 5'_3DR_WT_R | CTCAGCAGCTATTAAACCAGTAGG |
| 5'_3DR_KI_R | CACTCTGTTGTTGTCCCTATCG |
| 3'_3DR_Comm_R | ATATTCTCCACAGCCAGCGTG |
| 3'_3DR_WT_F | AGACAAGTATATACTCAAGGTGAGTCG |
| 3'_3DR_KI_F | CAAGATTTGCCAAAGCGCTTG |
| 3DR_CreChk_F | CGATAGGGACAACAACAGAGTGG |
| 3DR_CreChk_R | GCTTTGGCAAATCTTGATCTACAC |

D Genotyping results:

| Amplicon | Forward Primer | Position in WT | Position in KI, pre-Cre | Position in KI, post-Cre | Reverse Primer | Position in WT | Position in KI, pre-Cre | Position in KI, post-Cre | Size of Correct Amplicon (bp) | Result |
|---|---|---|---|---|---|---|---|---|---|---|
| 5' WT | 5'_3DR_Comm_F | 979 | 979 | - | 5'_3DR_WT_R | 2161 | - | - | 1182 | no amplicon |
| 5' KI | 5'_3DR_Comm_F | 979 | 979 | - | 5'_3DR_KI_R | 4387 | 1858 | - | 879 | 100% match by Sanger sequencing |
| 3' WT | 3'_3DR_WT_F | 1605 | 1605 | - | 3'_3DR_Comm_R | 2660 | 4886 | - | 1056 | no amplicon |
| 3' KI | 3'_3DR_KI_F | - | 4095 | - | 3'_3DR_Comm_R | - | 4886 | - | 792 | 100% match by Sanger sequencing |
| Cre Check | 3DR_CreChk_F | - | 1837 | 1837 | 3DR_CreChk_R | | 3863 | 2072 | 235 | 100% match by Sanger sequencing |

**Figure S2.** Overview of genotyping results confirming the successful introduction of 3 Cas12a DRs into the 3' UTR of murine *CTSD* via CRISPR-Cas. **A**, Genomic structure of the mouse *CTSD* gene prior to modification (top), after introduction of the "gene-trap" knockin (KI) allele (middle), and after removal of all elements besides the 3 Cas12a DRs using Cre-recombinase (bottom). The approximate positions of genotyping primers are indicated (blue arrows). **B**, Table showing the gRNA sequences using for CRISPR-Cas, the targeted endogenous sequence (with PAM sequence in red), the amino-acid sequences encoded by the sequence targeted by the gRNAs, and the modifications to gRNA-targeted regions within the KI targeting construct. **C**, Table showing the sequences for the DNA primers used for genotyping (with positions indicated in blue in panel **A** and identified numerically in panel **D**). **D**, Summary of genotyping results, showing the relative positions of individual primers, the predicted amplicon sizes for primer pairs, and the outcome of PCR amplification and subsequence Sanger sequencing. These results apply to a single clonal cell line chosen for use in downstream experiments. Note that no amplification was obtained for WT amplicons.

#### Supplementary Figure S3.

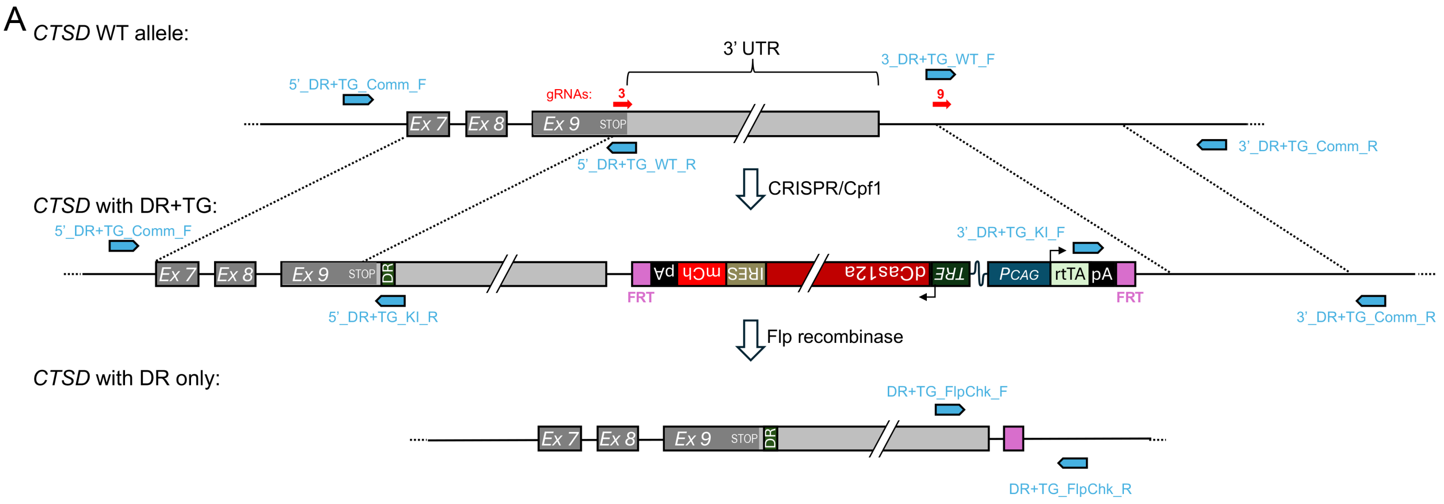


B gRNA design and alterations to knock-in insert:

| | gRNA_3 | gRNA_9 |
|---|---|---|
| **gRNA sequence** | **CCAATGCTGTCGTACTCTAAC** | **TGAATTGGGACCTTGGCGACG** |
| **Endog. sequence** | ...TTTGCCAATGCTGTCGTACTCTAACTT... | ...TTTCTGAATTGGGACCTTGGCGACG... |
| **A.A. sequence** | PheAlaAsnAlaValValLeuSTP | N/A |
| **KI insert sequence** | ...TTCGCGAACGCCGTAGTCCTGTGATAA... | deleted and replaced w/ FRT site |

C Genotyping primers:

| Primer Name | Primer Sequence |
|---|---|
| 5'_DR+TG_Comm_F | CTAGACAAGCTATGCCCTATACTCAC |
| 5'_DR+TG_WT_R | GTTAGAGTACGACAGCATTGGC |
| 5'_DR+TG_KI_R | GGAGCAAATCTACACTTAGTAGAAATTATTTTTTACTTATCAC |
| 3'_DR+TG_Comm_R | GATGCTTGTAGAACAACTGTAGTCC |
| 3'_DR+TG_WT_F | TTTCTGAATTGGGACCTTGGCG |
| 3'_DR+TG_KI_F | GATGCTTGTAGAACAACTGTAGTCC |
| DR+TG_FlpChk_F | CAGACAAGTATATACTCAAGGTGAGTCG |
| DR+TG_FlpChk_R | AGTTCTAGTTCAGACTCATCTTGCC |

D Genotyping results:

| Amplicon | Forward Primer | Position in WT | Position in KI, pre-Flp | Position in KI, post-Flp | Reverse Primer | Position in WT | Position in KI, pre-Flp | Position in KI, post-Flp | Size of Correct Amplicon (bp) | Result |
|---|---|---|---|---|---|---|---|---|---|---|
| 5' WT | 5'_DR+TG_Comm_F | 1219 | 1219 | - | 5'_DR+TG_WT_R | 1886 | - | - | 668 | no amplicon |
| 5' KI | 5'_DR+TG_Comm_F | 1219 | 1219 | - | 5'_DR+TG_KI_R | - | 1924 | - | 706 | 100% match by Sanger sequencing |
| 3' WT | 3'_DR+TG_WT_F | 2565 | - | - | 3'_DR+TG_Comm_R | 3218 | 12718 | - | 653 | no amplicon |
| 3' KI | 3'_DR+TG_KI_F | - | 11638 | - | 3'_DR+TG_Comm_R | 3218 | 12718 | - | 1080 | 100% match by Sanger sequencing |
| Flp Check | DR+TG_FlpChk_F | - | - | 2463 | DR+TG_FlpChk_R | - | - | 2836 | 374 | 100% match by Sanger sequencing |

**Figure S3.** Overview of genotyping results confirming the successful introduction of the DR+TG insert into the 3' region of murine *CTSD* via CRISPR-Cas. **A**, Genomic structure of the mouse *CTSD* gene prior to modification (top), after introduction of the DR+TG knockin (KI) allele (middle), and after removal of the TG portion using Flp-recombinase (bottom). The approximate positions of genotyping primers are indicated (blue arrows). **B**, Table showing the gRNA sequences, the targeted endogenous sequences (with PAM sequences in red), the amino-acid sequence encoded by the sequence targeted by gRNA_3, and the modifications to gRNA-targeted regions within the KI targeting construct. **C**, Table of DNA primers used for genotyping. **D**, Summary of genotyping results, showing the relative positions of individual primers, the predicted amplicon sizes for primer pairs, and the outcome of PCR amplification and subsequence Sanger sequencing. These results apply to a single clonal cell line chosen for use in downstream experiments. Note that no amplification was obtained for WT amplicons.
